# Propofol-Induced Unresponsiveness is Associated with Impaired Feedforward Connectivity in the Cortical Hierarchy

**DOI:** 10.1101/213504

**Authors:** Robert D. Sanders, Matthew I Banks, Matthieu Darracq, Rosalyn Moran, Jamie Sleigh, Olivia Gosseries, Vincent Bonhomme, Jean-François Brichant, Mario Rosanova, Aeyal Raz, Giulio Tononi, Marcello Massimini, Steven Laureys, Mélanie Boly

## Abstract

**Background:** Impaired consciousness has been associated with impaired cortical signal propagation following transcranial magnetic stimulation (TMS). Herein we hypothesized that the reduced current propagation under propofol-induced unresponsiveness is associated with changes in both feedforward and feedback connectivity across the cortical hierarchy.

**Methods:** Eight subjects underwent left occipital TMS coupled with high-density electroencephalograph (EEG) recordings during wakefulness and propofol-induced unconsciousness. Spectral analysis was applied to responses recorded from sensors overlying six hierarchical cortical sources involved in visual processing. Dynamic causal modelling (DCM) of evoked and induced source-space responses was used to investigate propofol’s effects on connectivity between regions.

**Results:** Propofol produced a wideband reduction in evoked power following TMS in five out of six electrodes. Bayesian Model Selection supported a DCM with hierarchical feedforward and feedback connections to best fit the data. DCM of induced responses revealed that the primary effect of propofol was impaired feedforward responses in cross frequency theta/alpha-gamma coupling and within frequency theta coupling (F contrast, Family Wise Error corrected p<0.05). An exploratory analysis (thresholded at uncorrected p<0.001) also suggested that propofol impaired feedforward and feedback beta band coupling. Posthoc analyses showed impairments in all feedforward connections and one feedback connection from parietal to occipital cortex. DCM of the evoked response potential showed impaired feedforward connectivity between left sided occipital and parietal cortex (T contrast p=0.004, Bonferroni corrected).

**Conclusions:** Our data suggest that propofol-induced loss of consciousness is associated with reduced evoked power and impaired hierarchical feedforward connectivity following occipital TMS.

## Introduction

Impaired consciousness has been associated with impaired cortical signal propagation and complexity of cortical responses to transcranial magnetic stimulation (TMS)^1–6^. It is hypothesized that these effects reflect impaired integration of information within the thalamo-cortical system^7–9^. In these terms, consciousness (defined as “subjective experience”) results from the continuous bidirectional flow of information between hierarchically organized thalamo-cortical units forming an anatomical “small world structure”^10^. In this paper we use the term “feedback” to refer to information moving down from higher-order association cortices to lower order primary sensory relay brain regions; “feedforward” is the opposite i.e. information moving in a same centripetal direction as the traditional sensory pathways. Accumulating data show that cortical connectivity is reduced under anaesthesia^11–19^, with increasing evidence from resting state data (collected in the absence of overt sensory stimuli) that feedback connectivity is predominantly suppressed (though see^20^). Moreover our recent study in rodents found direct evidence that feedback cortico-cortical connections are preferentially suppressed by isoflurane^21^. However these observations seem at apparent odds with the impaired current propagation observed across the cortex following TMS that would also recruit feedforward pathways, especially if targeted to a lower order cortical region. We hypothesized that recruiting both feedforward and feedback projections through TMS of sensory cortex may reveal impairment in bidirectional connectivity.

Introduced by Hubel and Wiesel^22,23^, the concept of a cortical hierarchy is proposed to underlie an ascending information processing stream, with lower-order sensory regions converging on increasingly complex multimodal association cortices. This has been further developed with models based on predictive coding^25,24^, which emphasize the role of descending connections in the integration of information across functionally specialized brain regions. Using TMS to stimulate a lower level of the hierarchy affords the opportunity to model bidirectional connectivity between different levels of the cortical hierarchy.

To provide a mechanistic account of these effects, we employed dynamic causal modelling (DCM) to investigate changes in connectivity across the cortical hierarchy in source space. Our primary outcome was DCM of the induced response as it allows modelling of cross-frequency coupling that is important for information transfer across the cortical hierarchy^24,26^. We previously used DCM to model resting state EEG data between two higher order cortical regions (finding impaired feedback connectivity)^18^; herein we extend these models to include an additional (lower) level of the cortical hierarchy. DCM offers several advantages for testing hypotheses about between- and within-region coupling, including the use of Bayesian Model Selection (BMS) to choose the most plausible model for the data^10,11^. Our primary outcome was to assess connectivity changes in the induced response induced by propofol. DCM of the induced response allows estimates of “between region”, “within frequency” and “cross frequency” coupling; all are important for the integration of information between hierarchical regions of cortex. Regarding the latter point, most studies of connectivity between different cortical regions under anesthesia have focussed on “within frequency” effects (e.g.^27,28^). DCM of induced responses models time-varying spectral changes and the interaction between these spectra at different sources. Essentially this provides assessment of power-based connectivity strengths between multiple sources over time, including the ability to look at cross-frequency power interactions (i.e. amplitude-amplitude coupling). It is important for the reader to note that this is different to other forms of cross-frequency coupling, that are also biologically important, and typically have been assessed using phase-phase or phase-amplitude coupling rather than power coupling^29–31,30,32–34^. We also leverage the DCM of the ERP, which includes priors about the neuronal circuitry, and fit the data with a biological model (DCM for the induced response does not employ such a biological model). In doing so, we sought to verify the changes that we observed in the DCM of the induced response with those of an alternate model.

## Materials and Methods

This study was approved by the Ethics Committee of the Medical School of the University of Liege. Following informed consent, eight subjects underwent occipital (BA19) TMS-EEG (~110V/m) during wakefulness with subjects lying on the bed with the eyes open. Following placement of an intravenous catheter, a target controlled infusion (TCI, Alaris TIVA, CareFusion) of propofol was commenced by a certified anesthesiologist. Propofol was infused until the subject became unresponsive to verbal command or mild shaking (Ramsay sedation scale 5 ^32^) and maintained at stable plasma concentration through TCI. Plasma and effect site concentrations of propofol were estimated using a three-compartment model^33^. TMS-EEG was then repeated in this unresponsive state. Throughout the experiment, oxygen was administered through a loosely fitted facemask. None of the subjects recalled events after recovery from propofol-induced unresponsiveness.

TMS was combined with a magnetic resonance-guided navigation system (NBS) and a 60-channel TMS-compatible EEG amplifier (Nexstim eXimia, Nexstim Plc, Finland). Real-time navigation based on individual structural magnetic resonance images was used to optimize the efficacy of TMS. The maximum electric field induced by TMS was always oriented perpendicularly to the convexity of the occipital cortical gyrus and its intensity was adjusted to values above the threshold for a significant EEG response (80-160 V/m). The reproducibility of the stimulation-coordinates across sessions was optimized by software coupled to the NBS system that indicated in real-time any deviation from the desired target greater than 3 mm. TMS was performed by means of a Focal Bipulse 8-Coil, driven by a Mobile Stimulator Unit (Eximia TMS Stimulator, Nexstim Plc.). At least 200 stimuli were acquired, with stimuli delivered at random intervals (between 2 and 2.3 s). Auditory response to the coil’s click and bone conduction were minimized as in previous studies^34^. Occipital TMS in wakefulness did not induce reports of changes in visual perception.

### Preprocessing

EEG data were filtered at 0.5-40 Hz (finite impulse response, EEGLAB). Higher frequencies were not analysed over concern of EMG contamination of the signal following TMS. Wakefulness and unresponsiveness data were then downsampled to 250 Hz, epoched (- 800ms to +800ms around TMS pulse), and referenced to average using the Statistical Parametric Mapping software (SPM12, <u>www.fil.ion.ucl.ac.uk/spm</u>). For descriptive purposes, we describe the data using the following bands: theta 4-8Hz, alpha 8-14Hz, beta 14-28Hz and gamma 28-40Hz.

Time-frequency analysis in sensor space was then conducted for 6 selected channels in occipital (PO3, PO4), parietal (CP3, CP4) and frontal (AF1, AF2) regions using a 7^th^ order Morlet wavelet transform followed by robust averaging. These regions were selected as overlying hierarchical-connected regions of cortex.

### Statistical Analysis of sensor level data

Group level contrasts were conducted at each channel for changes in the spectral response to TMS induced by propofol using T contrasts. Statistical significance was set a Family Wise Error rate corrected for multiple comparisons of p<0.05 of the peak response.

### Source reconstruction

Source reconstruction used a realistic boundary element method head model and equivalent current dipoles for DCM of the induced response^31,31^ and imaging model for evoked response potential^31,31^ centered on the coordinates of six cortical sources selected based on their hierarchical connectivity with the occipital lobe^36,36^. Source prior locations and coordinates were: left inferior occipital gyrus (MNI -27 -97 -10), right inferior occipital gyrus (27 -97 -10), left superior parietal lobule (-26 -64 56), right superior parietal lobule (26-64 56), left dorsolateral prefrontal cortex (-48 40 20) and right dorsolateral prefrontal cortex (48 40 20).

### Dynamic Causal Modelling

DCMs can be divided into two classes: biophysical and phenomenological^30^. Biophysical models (e.g. for the evoked response potential [ERP] or functional magnetic resonance imaging) include constraints imposed by the biology of the underlying neuronal circuits such as membrane properties of pyramidal cells or interneurons. In contrast, phenomenological models (e.g. for the induced response) describe the statistical relationships between factors in the model, for induced responses the critical factors are the power spectra themselves. Our primary interest was to model the statistical relationships in the power spectrum across cortex and hence we focus on the DCM of the induced response. We did this as cross frequency coupling is an important method of inter-regional information transfer in hierarchical regions of cortex. As a secondary analysis, that we hoped would confirm the findings for the induced response, we conducted DCM of the ERP^31^.

### Dynamic Causal Modeling of Induced Responses

For thorough description of DCM of induced responses see^30,31^. In brief, DCM of induced responses models the evolution of instantaneous power where temporal changes of power in a source are modelled as a network function of power in all sources. The equation for the model is pasted below, where *g*(*ω*)_*i*_ is the spectral density, over frequency *ω*, of the *i*-th unit.

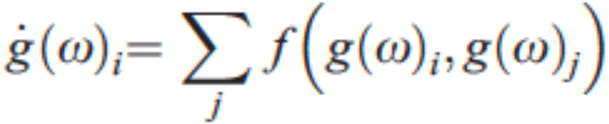

This model explains the relationship between the amplitude of an oscillation in one region influencing that in another. DCM utilizes a generalized convolution model of the coefficients of its Taylor expansion (the input) as the outputs. Causal inference is permitted as this is a “functional” expansion where time information is retained so we know that the relevant inputs precede the outputs. DCM of induced responses is therefore models time-dependent changes in spectral energy. We can therefore model the dynamics of the equation above suing a first order Taylor expansion to give:

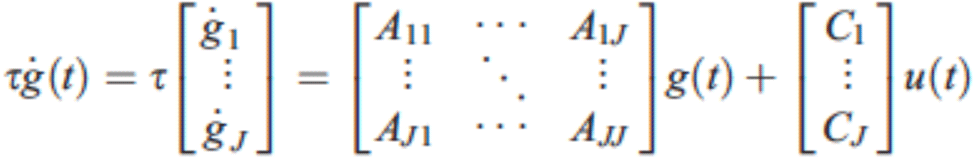

In this context the matrices A and C contain coupling parameters that explain changes in spectral activity based on spectral changes (either from other sources or within the same “self” source) and inputs (TMS), *u(t)*, to the model. A and C can be further defined:

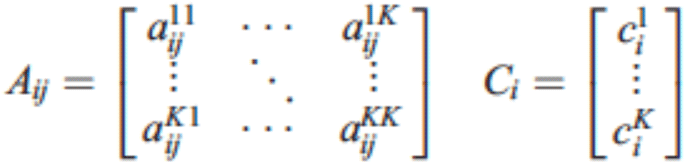

Such that the scalar a_ij_^kl^ relays how changes in the k^th^ frequency in the i^th^ source depend on the l^th^ frequency in the j^th^ source. Similarly c_i_^k^ explains the frequency-specific influence of the TMS (the input) on the k^th^ frequency of the i^th^ source. Together this allows modelling of “within frequency” and “cross frequency” coupling, “within (self modulation)” and “between” sources.

### Bayesian Model Selection

For the DCM analysis of induced responses, individual TMS-EEG trials were linearly detrended and Bayesian model selection (BMS) was used to optimize model parameters for the analysis, including optimal onset time for the DCM of cortical response to TMS (testing 1-20ms onset times post TMS to 400ms after TMS; data not shown), inclusion of “within region” intrinsic (self) modulation at a source, the modulation of ascending and/or descending connections, the presence of connectivity modulation at lower and/or higher cortical hierarchy levels (so called ‘B parameters’), and the presence of linear and/or non-linear (cross-frequency) effects. The model with highest posterior probability for each class of model parameter was selected for the final DCM models.

For the DCM of the ERP, BMS was used to optimise the optimal onset time of cortical response to TMS (testing 1-20ms post TMS), the modulation of ascending and/or descending connections and the presence of connectivity modulation at lower and/or higher cortical hierarchy levels as well as type of source reconstruction model used. The only difference in the parameters chosen by the BMS for DCM of the ERP compared to the induced responses was the source model used (Imaging not Equivalent Current Dipole).

### Statistical Analysis of DCM estimates for the induced response

Parameter estimates of changes in connectivity between wakefulness and propofol sedation (B parameters) obtained from the Bayes-optimal DCM model were entered in a full factorial design with eight levels (one per connection) and three factors: feedback versus feedforward directions, lower (occipital to parietal) versus higher (parietal to frontal) hierarchical levels, and left versus right hemispheres. A first omnibus F-contrast investigated for any effect of modulation of cortical connectivity between wakefulness and propofol sedation (using an eye(8) F-contrast as implemented in SPM). A further F-test searched for an effect of hierarchical level on connectivity changes. Finally, post-hoc t-tests investigated the presence of increased versus decreased cross-frequency coupling in feedforward or feedback connections in propofol compared to wakefulness: two contrasts used zeros for all feedback connections and 1 or -1 for feedforward connections, and two other contrasts used zeros for all feedforward connections and 1 or -1 for feedback connections. The final post hoc analyses used a -1 per connection and zeros for all other connections. All results were corrected for multiple comparisons across frequencies using peak-based Family Wise Error (FWE) p<0.05 of the peak response. Where appropriate further exploratory analyses were conducted with p<0.001 of the peak response.

### Statistical Analysis of DCM estimates for the ERP

BMS was used to confirm the same factors as in the DCM for the induced response however the model of source reconstruction was also tested. Contrast estimates for the change in connectivity induced by propofol (compared to wake) were obtained from the “B parameters” in the DCM. One sample T-tests were then conducted with a null hypothesis of zero change (i.e. mean values of 0). Statistical significance was set as a Bonferroni corrected p value of 0.05/8 = 0.00625 (multiple corrections for the eight connections tested).

## Results

### Propofol induces a loss of evoked power in EEG responses within single electrodes

A decrease in evoked power was observed in most electrodes during propofol compared to wakefulness (**Figure 1**). In particular, occipital channels PO3 and PO4 showed decreased power respectively at 11 Hz (peak level FWE p=0.0127) and 21 Hz (peak level FWE p=0.045). In parietal regions, channel CP3 showed decreased evoked power at 12 Hz (peak level FWE p=0.017) and 18 Hz (peak level FWE p=0.020) and channel CP4 showed decreased power at 29 Hz (peak level FWE p=0.034) and 33 Hz (peak level FWE p=0.044). Frontal channel AF1 also showed decreased 40 Hz evoked power (peak level FWE p=0.045). In summary, and consistent with prior data of the natural frequencies of different brain regions following TMS^38^, the peak level power differences observed in occipital and parietal cortex were in the alpha and beta frequencies, and a peak in the frontal cortex in gamma frequencies.

**Figure 1:**
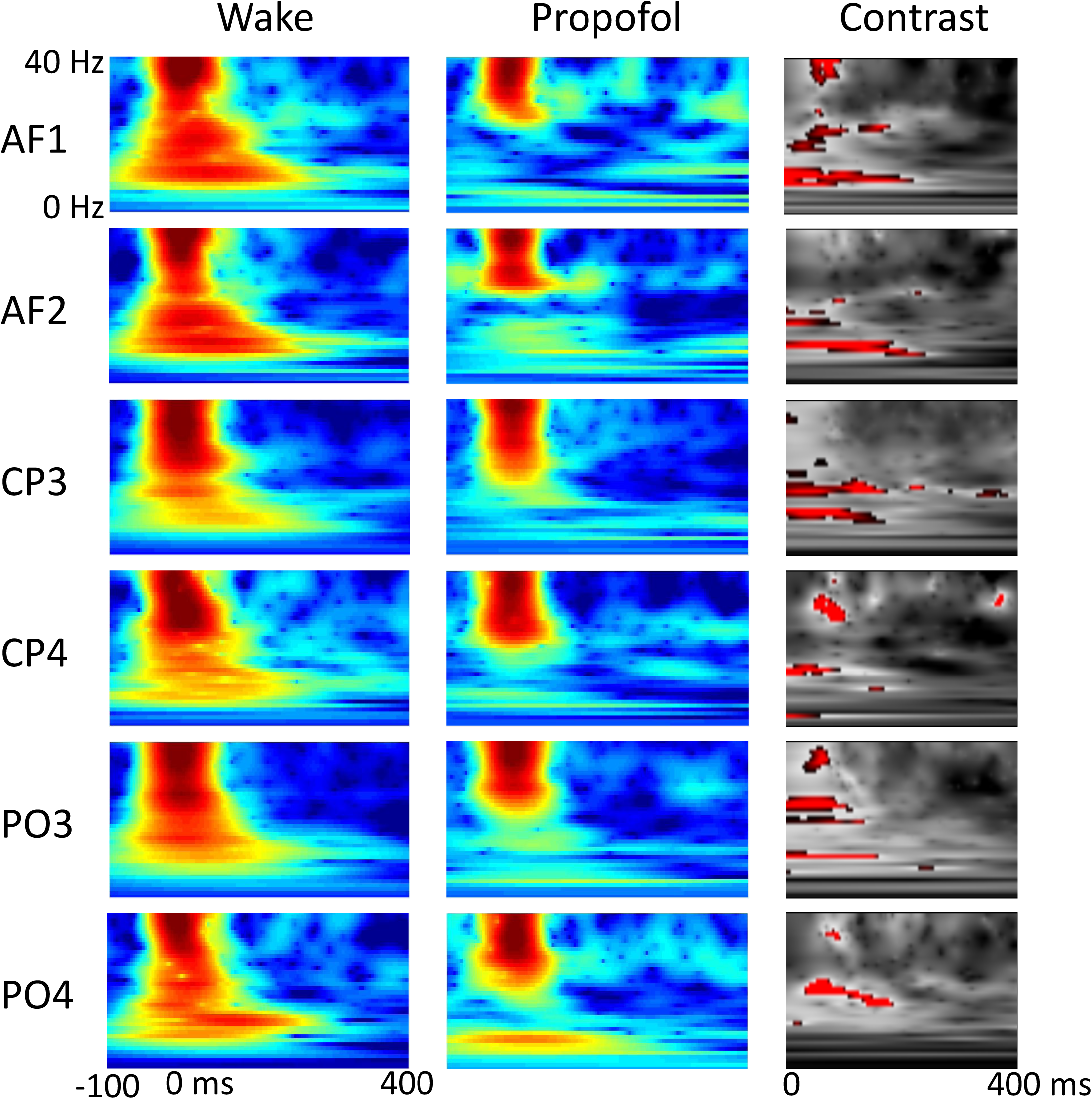
Sensor-space EEG responses to TMS in wakefulness and propofol: displayed for frontal electrodes AF1 and AF2, parietal electrodes CP3 and CP4 and occipital electrodes PO3 and PO4 shown in rows. The columns show the wake and propofol time frequency responses at the respective electrodes from -100 to 400ms after the TMS and in the last column the T contrast results for the propofol-induced decreases in TMS-EEG evoked power between 1-400ms. Red implies decreased power during propofol compared to wakefulness (thresholded at p<0.001 uncorrected for display purposes to show possible changes for each electrode).

### Bayesian Model Selection: DCM of induced responses

To investigate the optimal parameters to include in the DCM of the data Bayesian Model Selection was undertaken. Maximum model evidence was obtained for the following parameter choices: onset time of cortical responses = 1 ms (rather than 4, 8, 16 or 20 ms), intrinsic (self) modulation at each source, modulation of both ascending or descending connections rather than only one connection type (**Figure 2A**), and modulation of both occipito-parietal and fronto-parietal connections rather than only one cortical hierarchical level (**Figure 2B**). Critically, within the resulting model, including modulation of all hierarchical levels and connection types, modulating both linear and non-linear power connectivity was also given higher posterior evidence (**Figure 2C**). The optimal model is displayed in **Figure 2D**, with the connections numbered in the order used in subsequent figures.

**Figure 2:**
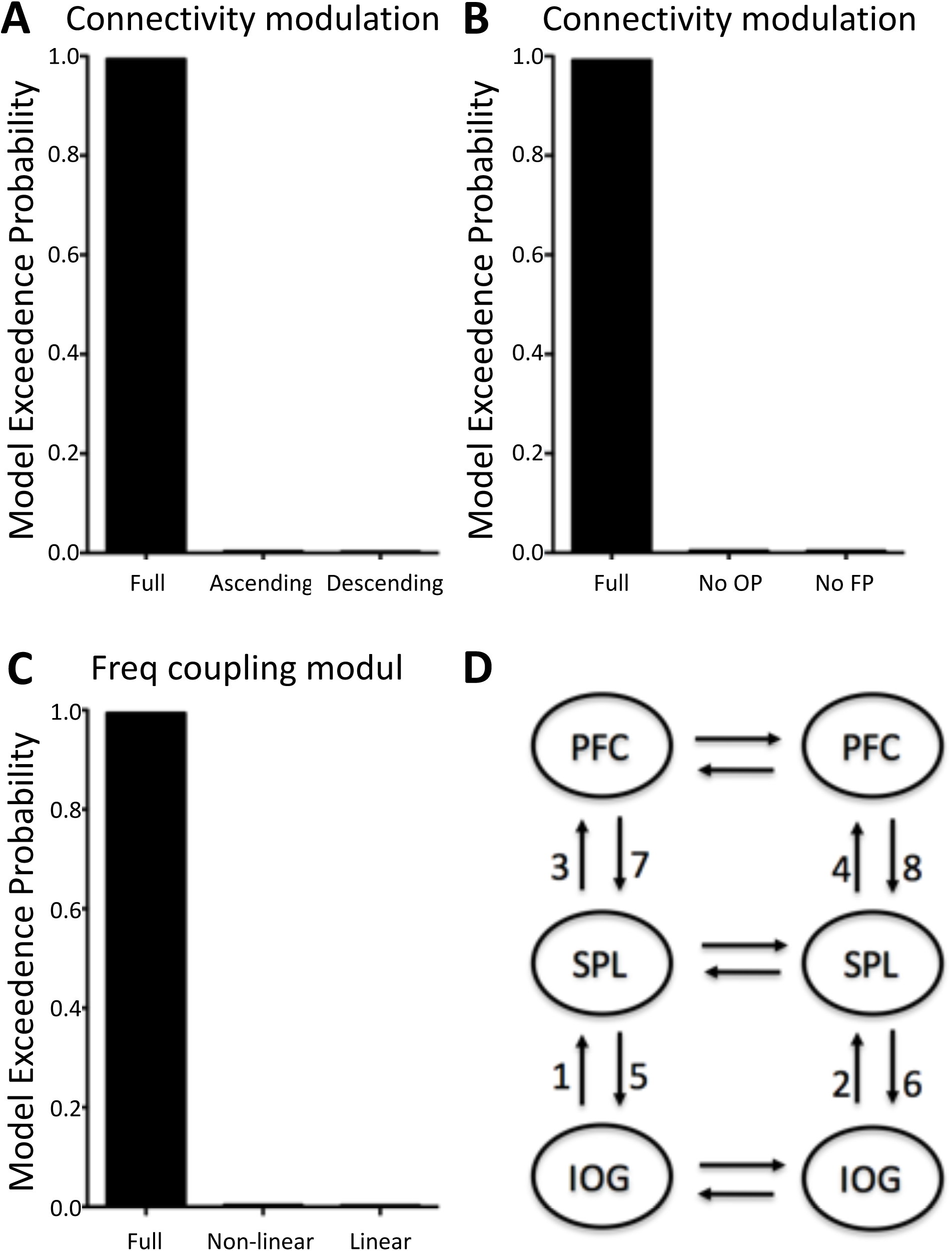
Bayesian model selection for the DCM models with different connectivity modulation profiles. (A) full model versus ascending connections only versus descending connections only and (B) full model versus full with no occipitoparietal connections versus full with no frontoparietal connections and (C) full model versus non-linear cross-frequency coupling (e.g. 8 Hz to 30 Hz) versus linear (e.g. 8 Hz to 8 Hz coupling) coupling modulations (B). The optimal DCM model selected after BMS is shown in (D). Numbers 1-8 in (D) refer to connection order for parameter estimates as displayed in Figures 3 and 4.

### DCM results: modelling the induced response shows reduced feedforward connectivity

An example of the wake and propofol source reconstructed and final DCM model data are shown in the **Supplementary Figure 1**. First, we used a full factorial design to analyse the connectivity changes predicted by DCM, including linear and non-linear frequency relationships, during propofol-induced unresponsiveness compared to wakefulness. In the DCM, linear connectivity relationships would occur between specific frequencies (e.g. 8 Hz to 8 Hz coupling) while non-linear connectivity refers to cross-frequency coupling (e.g. 8 Hz to 40 Hz). An omnibus F test (**Figure 3A**) revealed that propofol-induced unconsciousness was associated with altered theta to theta and theta/alpha to gamma coupling. In the theta band, coupling from peak frequencies at 8 to 8Hz (peak level FWE p=0.029; **Figure 3B**) was altered by propofol. Theta/alpha to gamma coupling from 8 to 40 Hz (peak level FWE p=0.008; **Figure 3C**) and 12 to 38 Hz (peak level FWE p=0.015; **Figure 3D**) was also reduced. These connectivity changes predominantly involved feedforward connectivity. A second peak, involving feedforward and feedback connectivity narrowly missed statistical significance (from 24 to 16 Hz, p<0.001 uncorrected corresponding to a peak level FWE p=0.097).

**Figure 3:**
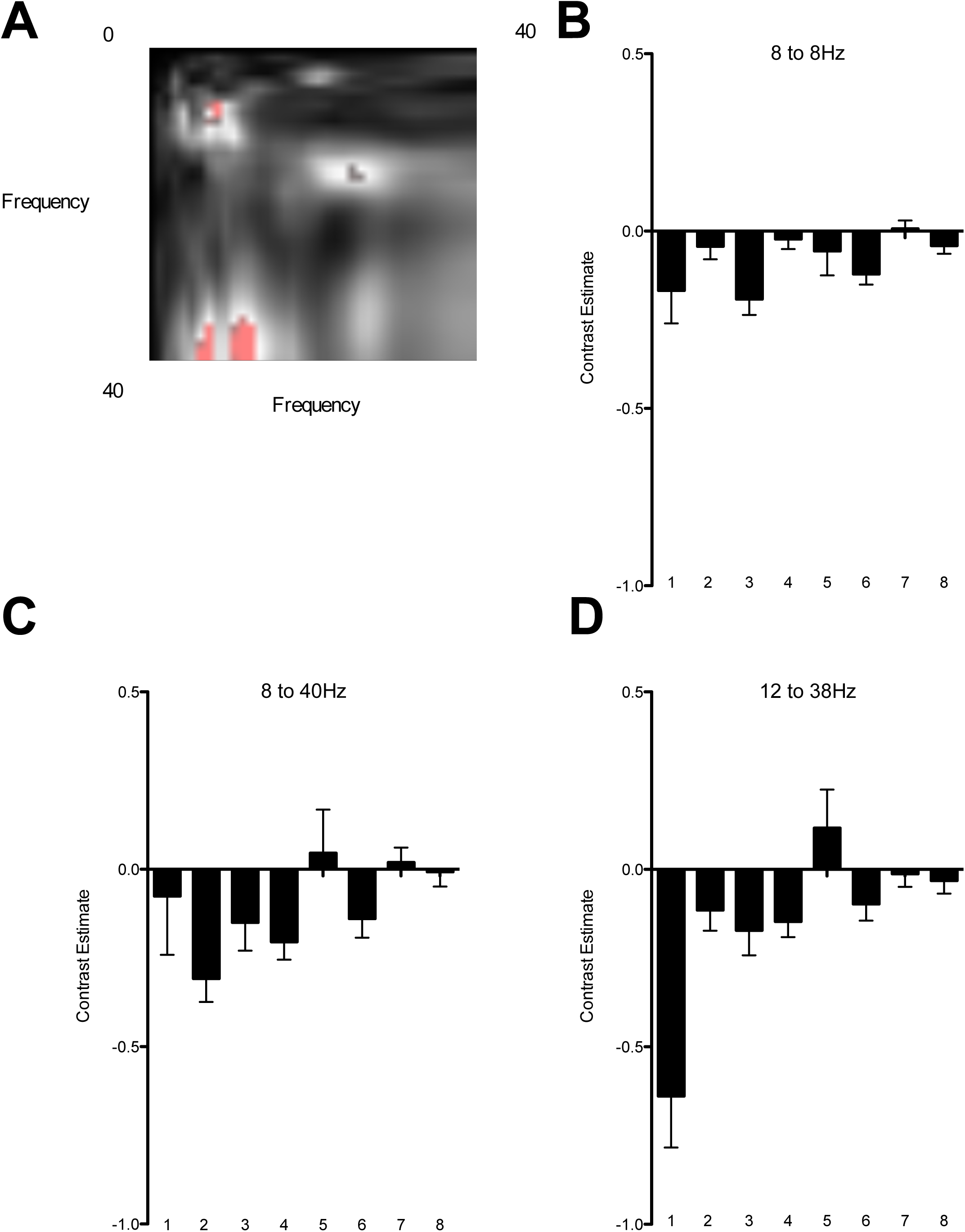
Decreased cross-frequency coupling throughout the cortical hierarchy during propofol-induced unconsciousness. (A) Frequency v. frequency plot displaying the significant clusters of altered cross-frequency coupling revealed by omnibus F-test in red. (B) Parameter estimates for change in connectivity strength 1 to 8 for propofol compared to wakefulness, for the maximum peaks of significance plotted in (A): 8 to 8 Hz (B), 8 to 40 Hz (C) and 12 to 38 Hz (FWE p<0.05).

### Post hoc Analyses

Subsequent post-hoc t-contrasts of the DCM data for induced responses, that analyse for frequency changes associated with feedforward or feedback processing, identified significant decreases in connectivity under propofol sedation compared to wakefulness in both feedforward and feedback directions. Feedforward connectivity was decreased in the alpha to gamma range (12 to 38 Hz; peak level FWE p=0.000). While feedback connectivity was decreased in the beta range (26 to 32 Hz; peak level FWE p=0.022 and 22 to 14 Hz; peak level FWE p=0.049) and in the alpha range (10 to 12 Hz; peak level FWE p=0.029). Though this model did not have the highest posterior evidence in Bayesian Model Selection, we confirmed similar findings with modeling from 20ms to 400ms, excluding any effect of the TMS artifact.

In order to identify which connections were involved in these changes we performed posthoc T contrasts over each connection (**Figure 4**). All feedforward connections showed depressed connectivity induced by propofol: left IOG to SPL (connection 1: 12 to 36Hz; peak level FWE p=0.009), right IOG to SPL (connection 2: 6 to 40 Hz; peak level FWE p=0.004), left SPL to PFC (connection 3: 8 to 8 Hz; peak level FWE p=0.018), right SPL to PFC (connection 3: 8 to 40 Hz; peak level FWE p=0.030). One feedback connection, right SPL to IOG, also exhibited depressed connectivity in various frequency bands (connection 6: 8 to 8Hz; peak level FWE p=0.014, 40 to 20Hz; peak level FWE p=0.031 and 26 to 36Hz; peak level FWE p=0.043).

**Figure 4:**
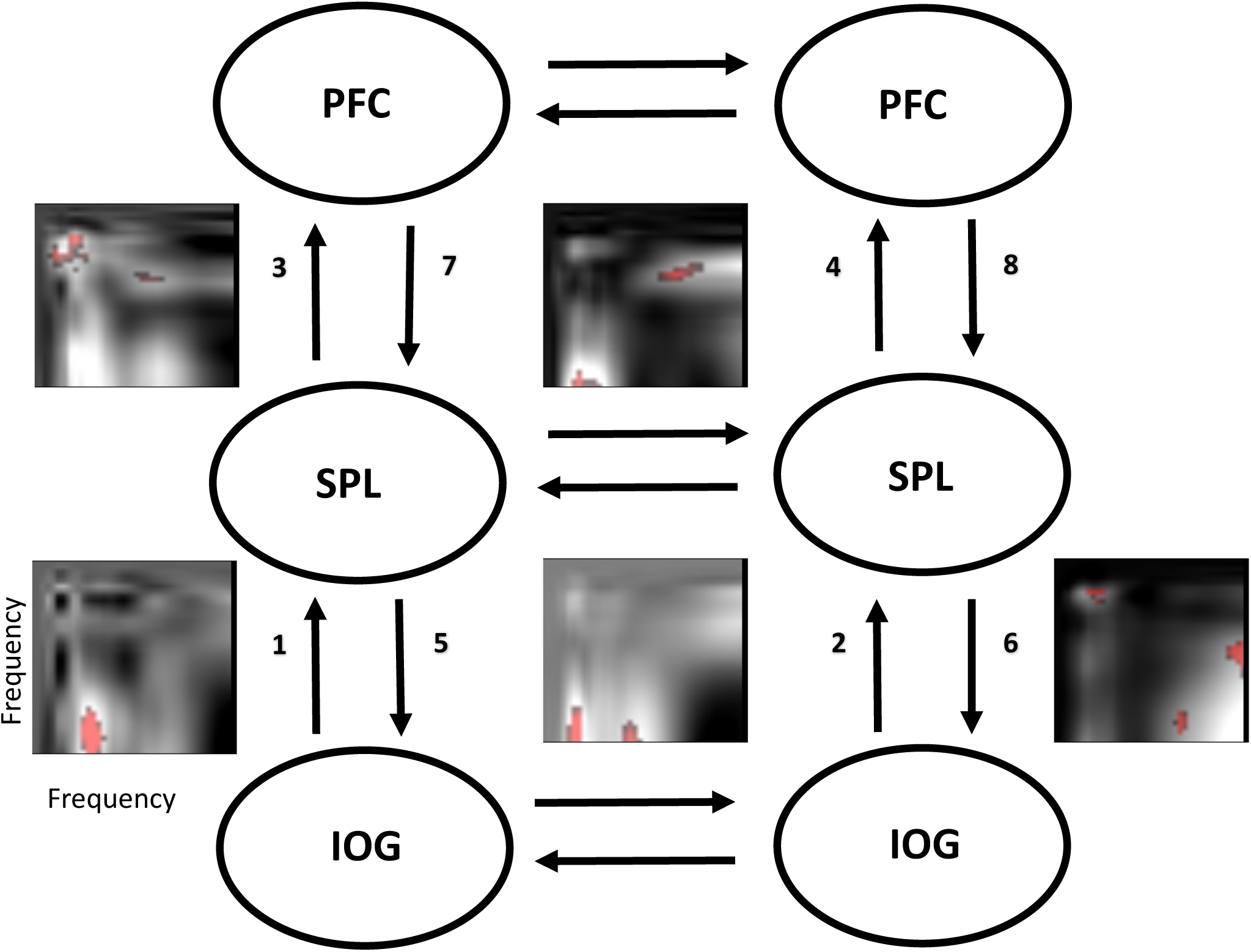
Post hoc T contrasts show that propofol impairs connectivity in five out of eight connections in DCM of the induced response. Across the 8 connections between inferior occipital gyrus (IOG), superior parietal lobule (SPL) and PFC (prefrontal cortex), five connections show significant differences in T contrasts (connections 1-4 and 6). For each connection where Family Wise Error corrected differences (p<0.05) were detected, a frequency-frequency plot is shown (to the left of the connection number for plots 1-4 that show feedforward connections) and to the right of connection number 6, a feedback connection). Red denotes significant impairments frequency coupling for that connection (FWE p<0.05).

### DCM results: modelling the evoked response potential shows reduced feedforward connectivity

BMS for the ERP model, showed that an Imaging model (with source optimization analogous to LORETA or minimum norm models^35^) was superior to an Equivalent Current Dipole approach (data not shown). An example of a single subject ERP is available in **Supplmentary Figure 2**. DCM of the ERP demonstrated a similar emphasis on the suppression of feedforward signaling during propofol-induced unresponsiveness (**Figure 5**). After Bonferroni correction for multiple comparison across eight connections, only the feedforward connection from left IOG to SPL showed significant impairment following left sided occipital TMS (p = 0.004). A similar result was obtained with modeling from 20 to 400ms following TMS, revealing the effected connection was feedforward left occipital to parietal (p=0.007).

**Figure 5:**
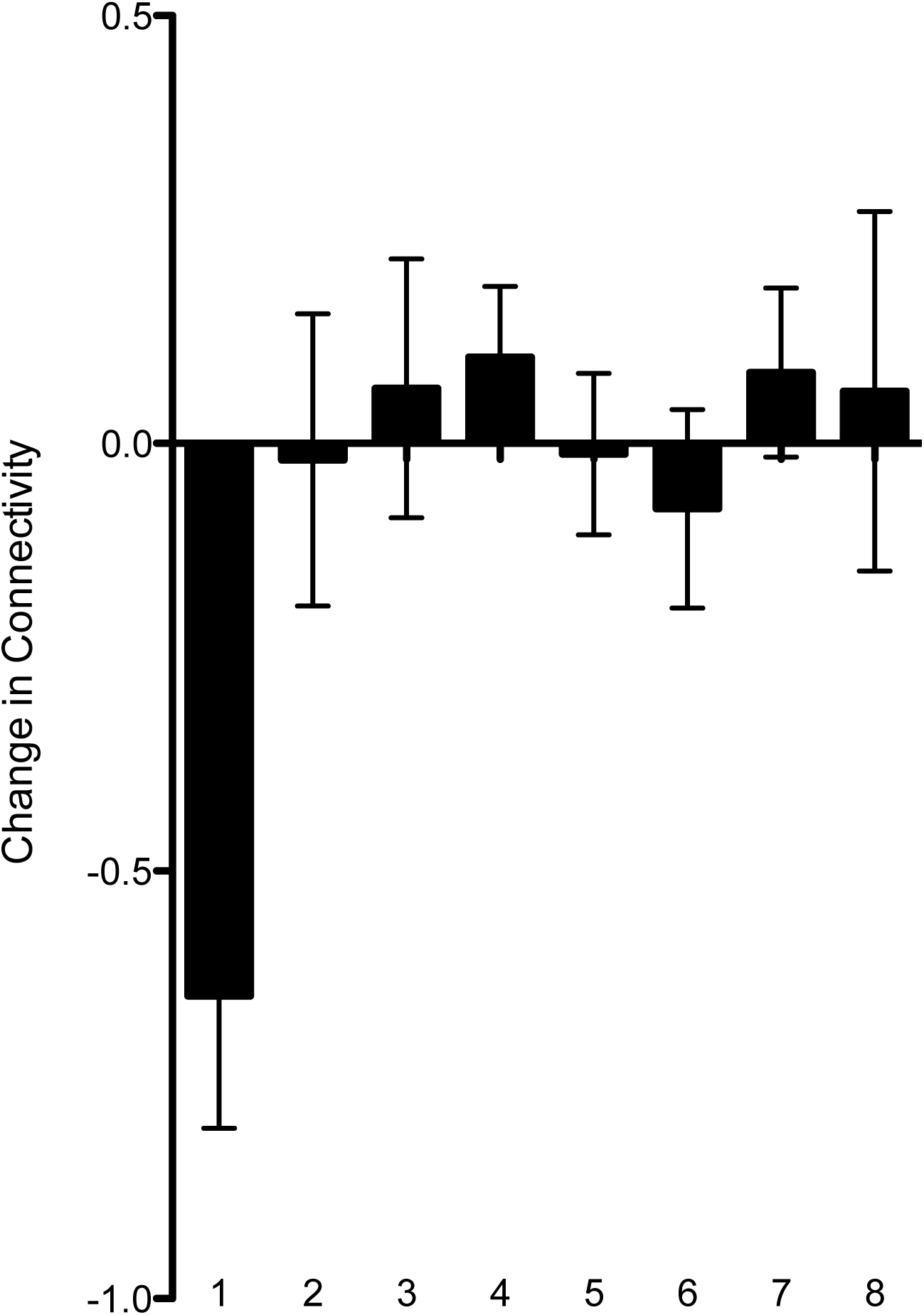
Propofol impairs feedforward coupling in a DCM of the ERP. Parameter estimates for change in connectivity strength for connections 1 to 8 for propofol compared to wakefulness. Connection 1 (feedforward from left IOG to left SPL) shows significantly impaired connectivity (Bonferroni corrected p<0.05).

## Discussion

Our data show that propofol diminishes evoked power following TMS across several sensors, with some variation in these sensors dependent on the underlying hierarchical regions of cortex. DCM of the induced power response and ERP following occipital TMS demonstrate impaired feedforward connectivity during propofol-induced unresponsiveness. Herein we formally described the diminished propagation of cortical responses following TMS targeted to a lower order cortical region^1,2,5^. This manifested as predominantly impaired feedforward connectivity, though feedback changes were also apparent in posthoc testing. This is important as the anaesthesia literature presently focusses heavily on feedback signalling. Our parsimonious explanation is that both feedback and feedforward connectivity is impaired under anesthesia, and each form is more easily revealed in different paradigms (resting state versus TMS evoked responses from lower order cortex). This is biologically plausible as resting state measures may emphasize feedback connectivity^18,28,39^, while evoked responses recruit feedforward pathways^24,40^. In this context our experiment was designed to identify bidirectional effects on connectivity in the cortical hierarchy.

Bayesian model selection of different DCMs suggested that the most complex model, i.e. one including both within and cross frequency coupling, and in feedback and feedforward directions, best explained our data. From this basis, we investigated the impact of propofol titrated to induce unresponsiveness, identifying predominantly an effect on feedforward connectivity. It may be argued that TMS, that is a direct cortical stimulus that bypasses any subcortical gate for signaling, is an artificial stimulus that lacks a biological simile. As such, the biological relevance of the stimulus may be questioned. Our counter argument is that TMS provides direct information about cortical processing^1,2,4,5^ and has been used successfully across cognitive neuroscience to this end. Furthermore TMS has recently shown remarkable diagnostic value to differentiate levels of consciousness^1^; we respectfully propose that understanding the cortical response to TMS will provide novel insights into the mechanisms of consciousness.

### Caveats

Of many methods of investigating connectivity, in this paper we chose to use DCM, based on our prior experience^18,41^ and our focus on modelling effects in the cortical hierarchy as explicitly as possible. All methods of connectivity assessment have potential pitfalls^42^ and DCM has been similarly critiqued^30,31,43,44^. Because DCM of the induced response does not rely on a specific biological model, it is not vulnerable to criticism (or praise) over the specifics of the biological model employed (unlike DCM for the ERP^30,31^). Furthermore its time-dependence, ensuring that inputs precede outputs, provides a similar to strength to the “gold standard” Granger Causality^42^. A strength over Granger Causality analysisis that DCM of induced responses permits inferences about cross-frequency coupling, which are prominent in cortical dynamics^29^. Nonetheless, DCM has weaknesses. It is computationally intensive, even when limited to eight frequency modes. Hence despite the apparent complexity of our modelling approach, DCM may overlook even more complex – but biologically important – relationships. The primary strength in our approach is to provide a model driven assessment of connectivity “between regions” and “cross frequencies”. This represents a novel approach for EEG analysis in the anesthesia literature which typically employs “within frequency” connectivity measures for effects “between regions”.

A further strength of our analysis is the robust statistical approach using family wise error correction based on Random Field theory to reduce type 1 error. We then supported our findings with the DCM for the ERP which employs a biological model, based on the canonical cortical microcircuit^24^. These models have proven very useful for probing conscious and unconscious states^41^, and herein the use of this biological model, confirms the effect on feedforward processing uncovered by DCM for the induced response. It is also important to note that the DCM for ERP (compared to the induced model) invoked a different source reconstruction technique providing further evidence that the DCM for the induced response produced a reproducible finding. Nonetheless we recommend that further studies are conducted in other subjects to confirm that our findings are truly reproducible.

While the focus of our study was to identify whether propofol interfered with connectivity including cross-frequency coupling, our scalp EEG recordings are not well suited to detect effects on ascending connectivity within higher gamma frequencies^45^. Future studies should use alternate methodologies, for example magnetonencephalography or intracranial recordings, to assess the role of high gamma activity. Also investigating the role of thalamus, while of potential theoretical interest^46,47^ was not technically feasible in the present study, but could be investigated in future studies particularly if using invasive recordings.

### Implications for the understanding of the impairment of consciousness induced by anesthesia

From a practical perspective, we have shown reduced power responses following TMS at several cortical electrodes. If this is shown in other paradigms to track levels of consciousness, a monitor may be derived that looks at the reduced power responses at a single electrode. This hypothesis will also need testing in further volunteers before clinical trial testing should begin. More fundamentally we show that feedforward signaling is impaired under anesthesia. Critically this involved theta-theta and theta/alpha-gamma coupling which prior studies have suggested are critical to feedforward processing^24,40,42,48^. These findings complement prior work showing that propofol impairs long-range cortical connectivity^18,49,50^ with feedforward and feedback connections being affected in different frequency bands^40,45,48,51^. While anesthetics are thought to affect cross-frequency coupling within region^52,53^ and phase-amplitude coupling within frontal and parietal electrodes^18,27,50,52,54^, our results suggest that cross-frequency power coupling *between levels of the cortical hierarchy* is affected by anesthesia. Theoretically this would affect integration of information across the cortical hierarchy.

Our experiment was designed to identify whether effects on feedforward processing could be detected following TMS, by targeting a lower order cortical region. As such we maximized sensitivity to identify this feedforward effect. This may have meant that feedback effects only came to prominence after lowering the statistical threshold on the F test for induced responses or in the posthoc T-tests. However we did not identify a strong signal on feedback processing in our DCM of the ERP. Nonethless, we find it plausible that propofol effects both feedforward and feedback processing and the next stage is to understand the convergence or divergence of these effects during clinical anaesthesia in the operating room. It is worth emphasizing that in our experiment feedback frontoparietal connectivity was not found to be affected by propofol unlike our prior DCM study^18^. Rather feedback SPL to IOG connectivity was impaired. This may relate to the specifics of our experimental paradigm (TMS targeted to the lower order cortex) and this should be tested through TMS applied to higher order regions to more directly recruit feedback frontoparietal connectivity.

A recent study showed that approximately 4.6% of patients may be aware of sensory stimuli (“connected consciousness”) under general anaesthesia, which is an important clinical problem^47,55^ and must involve ascending transmission of sensory information. As sensory cortices are still readily activated by surgical stimulation under anaesthesia^47^, and that connected consciousness can occur under anesthesia without frontal cortical activation^56^, future studies should assess how anesthesia effects feedforward connectivity following evoked sensory stimuli and what levels of the cortical hierarchy are involved. Further experiments recording TMS-EEG during states of both unconsciousness and of connected/disconnected consciousness should investigate if loss of hierarchical cross-frequency coupling within the cortex also contributes to anaesthesia-induced sensory disconnection.

## Conclusions

Our results suggest that cortical time-frequency spectral responses to TMS are perturbed by anesthesia. Furthermore impaired feedforward connectivity, employing cross-frequency coupling between hierarchical cortical regions, is evident during propofol-induced unconsciousness. These results shed light on the neural mechanisms of the loss of integration induced by propofol sedation in the cerebral cortex, and suggest that changes in both feedforward and feedback connectivity throughout the cortical hierarchy might be involved in anaesthesia’s effect on consciousness.

## Contributions

RDS, MD, JS, RM, AR, MIB Designed research, Analyzed data and Wrote the paper. MD, Analyzed data and Wrote the paper. OG, MAB, VB, JFB, MR, MM, SL Performed Research and Wrote the paper. GT Wrote the paper. MB Designed research, Performed Research, Analyzed data and Wrote the paper.

## Acknowledgments

OG a post-doctoral fellow, SL a research director at the national fund for scientific research (FNRS). This research was supported by the Belgian National Funds for Scientific Research (FNRS), Human Brain Project (EU-H2020-FETFLAGSHIP-HBP-SGA1-GA720270), Luminous project (EU-H2020-FETOPEN-GA686764), the European Commission, the James McDonnel Foundation, the Mind Science Foundation, the French Speaking Community Concerted Research Action (ARC-06/11-340), and the University and University Hospital of Liège. RDS, MD, AR and MIB are supported by the Department of Anesthesiology, University of Wisconsin, Madison. MIB additionally is supported by R01 GM109086. MR is partially supported by the grant Grant "Giovani Ricercatori" GR-2011-02352031 from the Italian Ministry of Health. MM is supported by James S. McDonnell Foundation Scholar Award 2013 and EU grant H2020 grant agreement 720270-Human Brain Project SGA1. MB is supported by R03NS096379.

---

**Supplementary Figure S1:**
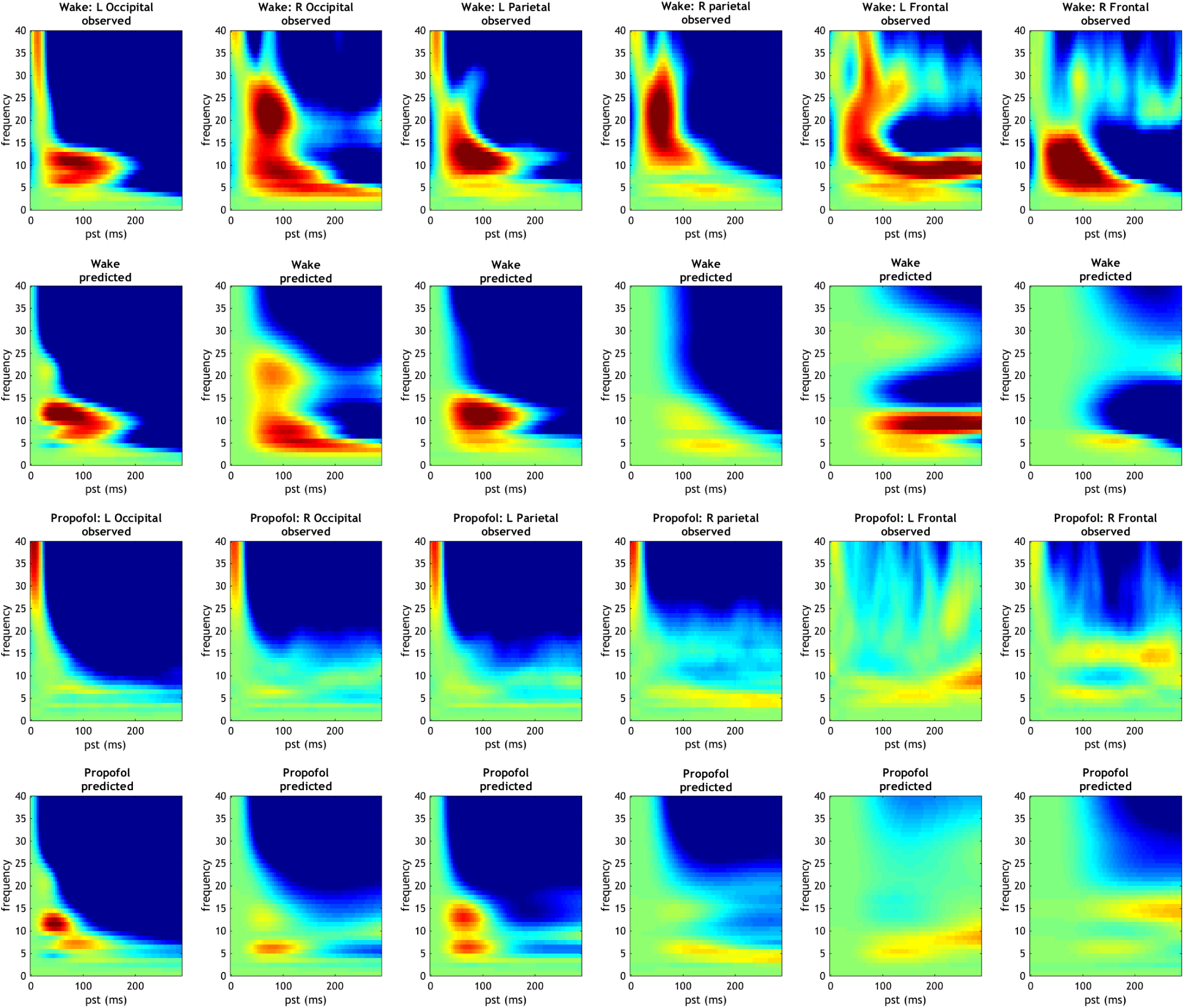
Example of source reconstructed time frequency (observed) and DCM of the induced response (predicted) of occipital TMS in wake and propofol-induced unresponsiveness from a single subject. Columns refer to left occipital, right occipital, left parietal, right parietal, left frontal and right frontal regions respectively.

**Supplementary Figure S2:**
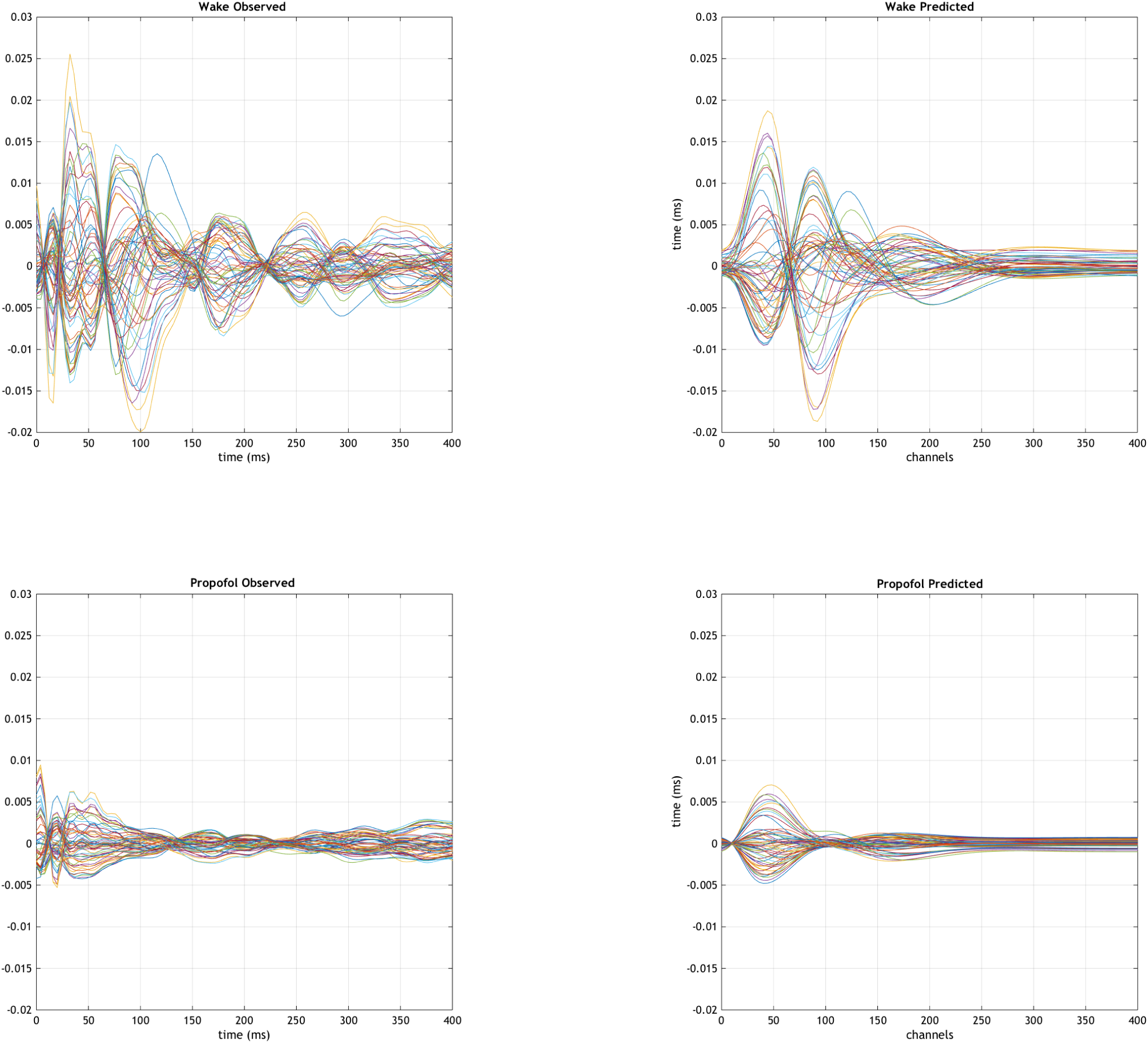
Example of source reconstructed ERP (observed) and DCM of the ERP (predicted) of occipital TMS in wake and propofol-induced unresponsiveness from a single subject. Columns refer to left occipital, right occipital, left parietal, right parietal, left frontal and right frontal regions respectively.

